# PTX3 Governs Fibroblast-Epithelial Dynamics in Lung Injury and Repair

**DOI:** 10.1101/2024.12.03.626584

**Authors:** Marie-Therese Bammert, Ines Kollak, Jan Hoffmann, Eva Peter, Holger Schlüter, Jun Li, Alexandre R. Campos, Coralie Viollet, Florian Gantner, Muriel Lizé, Matthew J. Thomas, Huy Q. Le

## Abstract

Dysfunctional interactions between fibroblasts and epithelial cells contribute to the progression of chronic lung diseases, including idiopathic pulmonary fibrosis (IPF). In this study, we utilized a coculture model of human small airway epithelial cells and fibroblasts to investigate intercellular communication during disease progression. Our transcriptomic and proteomic profiling reveal that fibroblasts repair epithelial cells in acute injury by boosting epithelial fatty acid metabolism; conversely, they exacerbate epithelial damage in chronic injury scenarios. By delineating regulators involved in these responses, we identified pentraxin 3 (PTX3) as a key antifibrotic factor secreted by fibroblasts in response to acute epithelial injury. Importantly, PTX3 levels are decreased in bronchoalveolar lavage (BAL) samples from IPF patients compared to non-fibrotic controls, indicating a potential link between diminished PTX3 levels and fibrosis progression. Furthermore, adding PTX3 to chronically injured epithelial-fibroblast cocultures mitigated the pro-fibrotic response and restored the epithelial barrier integrity. These findings highlight the dual roles of fibroblasts and the critical function of PTX3 in lung injury and repair, offering insights for therapeutic strategies.

## Introduction

Sensing and responding to changes in the microenvironment are key mechanisms of cells to ensure homeostasis within a tissue^1^. In response to extrinsic stimuli, cells adapt their cellular programs and transmit the information to the nucleus to fine-tune gene expression. Conversely, they secrete adjusted signals back in their environment. These processes rely on the coordination of various cellular synergies including canonical, mechanotransducing and metabolic sensing pathways. Thus, they are not restricted to individual cell but rather taking place in a complex network of cell-cell communication, fundamentally influencing cell fate decisions.^1–5^

Idiopathic pulmonary fibrosis (IPF) is a rare, devastating form of fibrosis in the lung, where excessive accumulation of extracellular matrix components occurs, which ultimately leads to distortion and increased stiffness of the lung architecture.^6^ Patients suffering from IPF have a poor prognosis with no regenerative therapies available. Whilst concrete initiating factors of IPF remain unclear, it is evident that chronic injury and aberrant epithelial repair, driven by disrupted communication between epithelial cells and fibroblasts, play a major role in the development of this fatal disease^7–9^. In contrast, acute lung epithelial injuries, such as those seen in mild lung diseases, are often temporary and can resolve over time with the support from surrounding cells, particularly through epithelial-fibroblast interactions^10–12^. However, unresolved injuries in severe cases impair proper intercellular communication and lead to long-term complications, including IPF^13–16^. This raises the question of why chronic injury persists in the lung, ultimately leading to fibrosis, while acute lung injury is temporary and resolves over time. Understanding these differences and how cells respond and alter their intercellular communication during tissue repair is essential for developing effective therapeutic strategies.

To address these questions, we established an air-liquid-interface model of chronically and acutely injured human small airway epithelial cells cocultured with lung fibroblasts. This model allowed us to identify key differences in the cellular response, focusing on the intercellular communication during acute and chronic injury. Our findings highlight the beneficial role of epithelial-fibroblast interaction in acute injury and elucidate the key mechanisms that facilitate effective wound healing. Notably, we discovered a protective effect associated with a metabolic shift towards fatty acid metabolism in epithelial cells, which appears to reinforce vital cellular processes and enhance resilience to injury. Intriguingly, secretomics analysis revealed that PTX3 plays a protective role in chronic injuries by mitigating the pro-fibrotic effect on epithelial cells. These findings provide valuable insights into the therapeutic potential of modifying the cellular niche within chronic injury conditions, to resolve fibrotic conditions within the lung.

## Results

### Chronic injury model exhibits IPF hallmarks

To explore the interplay between epithelial cells and fibroblasts in the human lung following injury, we developed a coculture system utilizing mature, differentiated primary small airway epithelial cells (SAECs) in an air-liquid interface and primary human lung fibroblasts (LFs). We investigated in two distinct scenarios that mimic chronic and acute injuries, each characterized by varying exposure times to TGF-β1, a known injury-inducing stimulus^17^. In the chronic injury model, SAECs pre-treated with TGF-β1 for 14 days (tSAECs) were employed, which were subsequently co-cultured with LFs or as monocultures for an additional 3 days in the presence of the stimulus (Fig. 1A). Conversely, no prior injury was induced in the acute model (nSAECs), but cells were cultivated, before the initiation of the coculture and TGF-β1 treatment.

**Figure 1:**
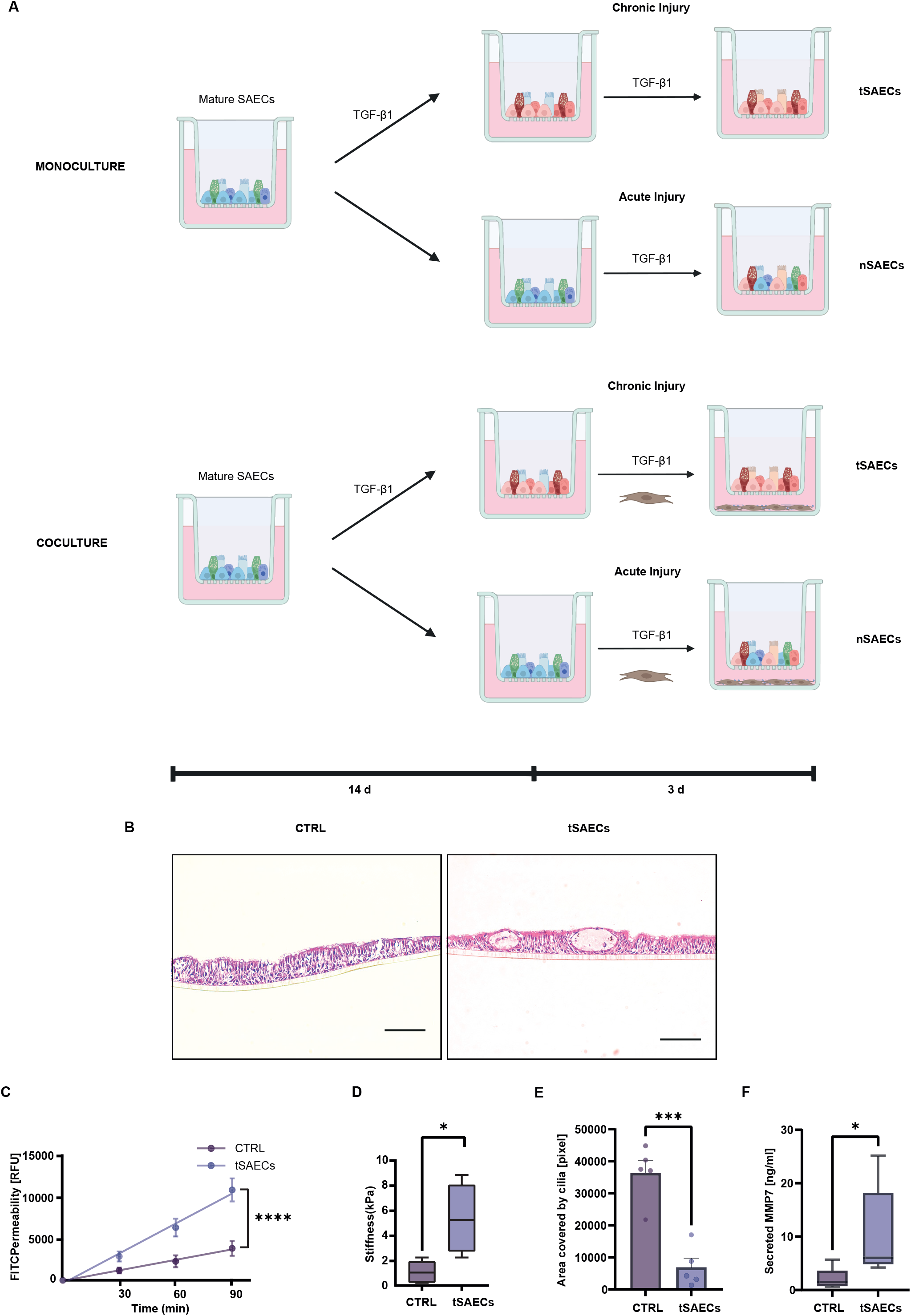
tSAECs recapitulate pro-fibrotic phenotype. **A,** Schematic overview of the cultivation and treatment strategy of nSAECs (acute treatment) and tSAECs (chronic treatment) in mono- and cocultures with fibroblasts. Mature SAECs (differentiated for 10 days) were exposed to TGF-β1 for 14 days and cultivated +/- fibroblasts and stimuli for another 3 days in a chronic injury setting. For the acute injury model, mature SAECs were differentiated further for 14 days without TGF-β1, before starting the coculture or TGF-β1 treatment. **B**, Representative H&E stainings of control (non-treated) and tSAECs showed damage of the epithelial barrier in tSAECs. **C**, FITC permeability measurement shows increased leakage of the epithelial barrier in tSAECs compared to control (n = 5, mean + s.e.m., t-test, ****p < 0.0001). **D**, Nanoindentation measurement shows increased viscoelasticity (Young modulus) in tSAECs compared to control (n = 5, mean + s.e.m., t-test, *p < 0.05) **E**, Quantification of ciliated cell area shows a significant decrease of active cilia in tSAECs compared to control (n = 5, mean + s.e.m., t-test, ***p < 0.001) **F**, ELISA measurement showed increase in biomarker MMP7 in tSAECs compared to control supernatants (n = 5, mean + s.e.m., t-test/Mann-Whitney, *p < 0.05).

To assess whether chronic exposure to TGF-β1 induces sustained injury and subsequent fibrosis-related alterations in the SAECs, we examined their morphological, physical, and integrity properties. Hematoxylin and eosin (H&E) staining unveiled pronounced structural differences and damage to the epithelial layer in tSAECs compared to control, untreated SAECs (Fig. 1B). Moreover, chronic stimulus exposure led to increased leakage of the epithelial barrier (Fig. 1C), reflecting the characteristic epithelial barrier damage and integrity loss observed in chronic lung diseases, including IPF^18^. This was accompanied by a significant increase in cellular stiffness in tSAECs compared to control, untreated SAECs (Fig. 1D), reflecting the increased rigidity of the IPF tissue and corroborating with measurements from IPF lungs^19^. Upon chronic treatment, tSAECs showed a decline in ciliated cells (Fig. 1E) as consistent with previous observations^20^, while cilia beating frequency remained unchanged (Supplementary Fig. S1A). Investigations in the secretion of pro-fibrotic factors revealed elevated levels of the clinical biomarker matrix metalloproteinase 7 (MMP7)^21^ (Fig. 1F) as well as MMP10^22^ (Supplementary Fig. S1B).

Taken this together, exposure to TGF-β1 for 14 days successfully induced a chronic injury and pro-fibrotic phenotype in SAECs, which resembles several key characteristics of IPF including increased stiffness, loss of the epithelial barrier integrity, decline in ciliated cells, and the secretion of biomarker MMP7.

### Acute and chronic injury on epithelial cells show distinct molecular responses

To investigate the molecular mechanisms distinguishing chronic and acute injury in epithelial cells, we performed RNA-seq on nSAECs and tSAECs. Our analysis of the gene expression profile in the monocultures revealed distinct pro-fibrotic and pro-inflammatory transcriptional responses in nSAECs and tSAECs compared to healthy controls (differentially expressed genes (DEGs), p-value < 0.05, Fig. 2A, B, Supplementary table S1A, S1B). Notably, tSAECs exhibited an increased number of transcriptional variances (total: 1422 DEGs) compared to the relatively modest changes observed in acute injury (total: 204 DEGs), reflecting the variable duration of injury exposure. Next, to identify how these global changes resemble human lung disease, clinical data from IPF patients^23,24^ was employed. Intriguingly, gene set enrichment analysis^25^ (GSEA) revealed a significant representation of gene signatures associated with central IPF tissue in tSAECs (enriched genes: 130; FDR: 0.00) highlighting their pronounced pro-fibrotic and -inflammatory nature, while nSAECs (enriched genes: 16; FDR: 0.02) lacked resemblance to IPF gene expression patterns (Fig. 2C, Supplementary Table S1C, S1D). Furthermore, disease-related analysis of upregulated genes in nSAECs showed significant involvement of genes implicated in lung infections (Supplementary Figure S2A, Supplementary Table S1E). This indicates that while mild pro-fibrotic responses present in early lung injury situations are detectable in nSAECs, the overall fibrotic gene signature compared to tSAECs is absent, making this a suitable tool to investigate key differences between chronic and acute epithelial injury.

**Figure 2:**
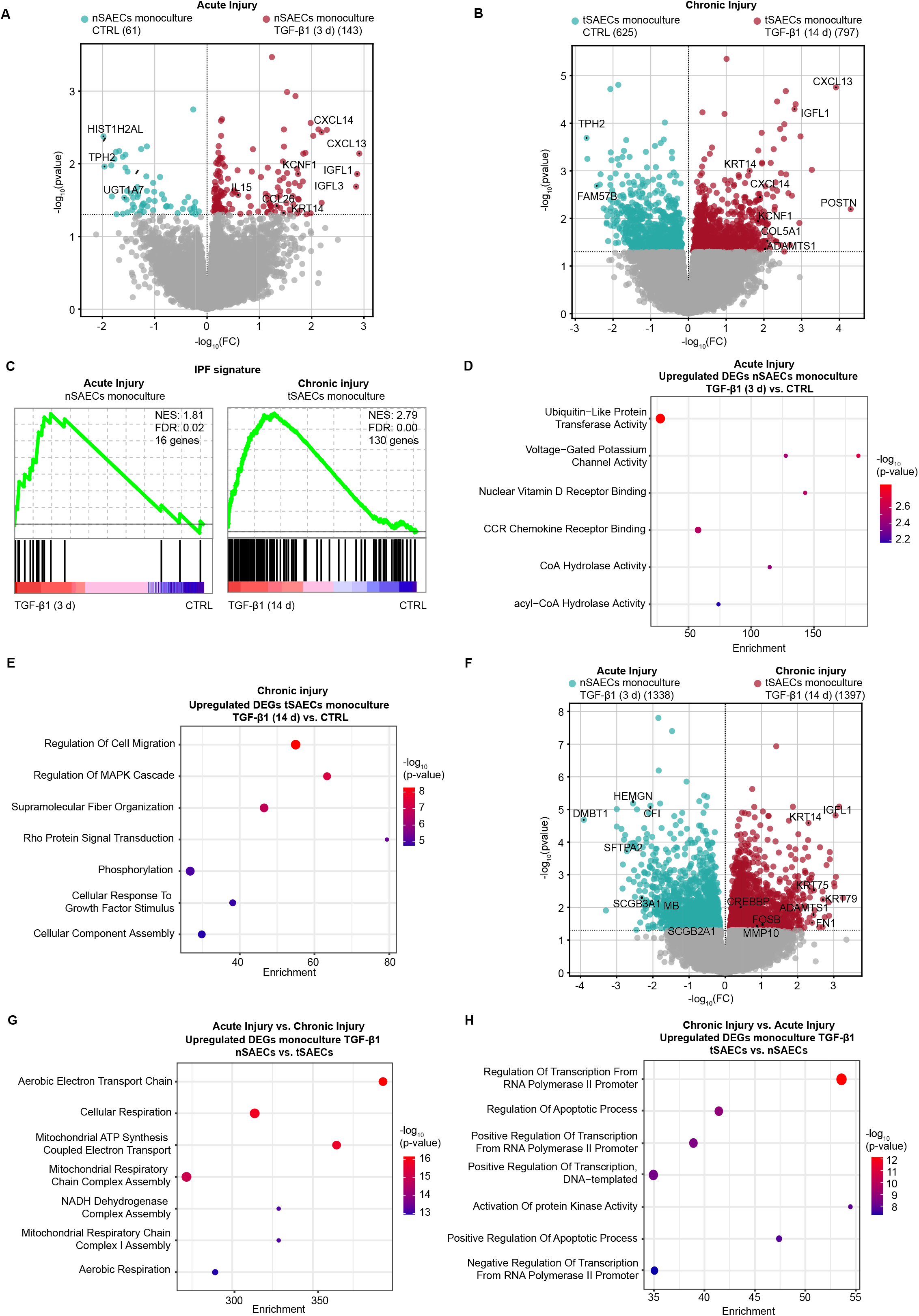
tSAECs and nSAECs exhibit different transcriptional programs depending on the injury. **A,** Volcano plot of all differentially expressed genes (DEGs) of nSAECs in monoculture comparing TGF-β1 treatment (3 d) vs. control (p-value < 0.05) shows differences in the transcriptome. **B**, Volcano plot of all DEGs of tSAECs in monoculture comparing TGF-β1 treatment (14 d) vs. control (p-value < 0.05) represents transcriptional differences among the conditions. **C**, Gene set enrichment analysis (GSEA) shows weak enrichment of an IPF lung gene signature in nSAEC monocultures comparing TGF-β1 treatment vs. CTRL, whilst tSAECs monocultures (TGF-β1 vs. CTRL) exhibit strong correlation for the IPF signature (p < 0.05). **D**, Gene ontology (GO) biological process analysis reveals increase in cellular stress responses in nSAECs TGF-β1 treated vs. CTRL (p < 0.05, log2 fold change (log2FC) > 0). **E**, Gene ontology (GO) biological process analysis shows increase in epithelial-to- mesenchymal transition (EMT) processes in tSAECs TGF-β1 treated vs. CTRL (p < 0.05, log2FC > 0). **F**, Volcano plot of all DEGs of tSAECs vs. nSAECs in monocultures with TGF-β1 (p < 0.05) shows differences in the expressed genes. **G**, Gene ontology (GO) biological process analysis shows increase in aerobic metabolism in nSAECs compared to tSAECs in monocultures with TGF-β1 (p < 0.05, log2FC > 0). **H**, Gene ontology (GO) biological process analysis shows increase in transcriptional activity in tSAECs compared to nSAECs in monocultures with TGF-β1 (p < 0.05, log2FC > 0).

Gene ontology (GO) enrichment analysis showed the induction of certain cellular stress responses in nSAECs, including membrane depolarization via potassium channels (Fig. 2D, Supplementary Table S1F), and changes in intracellular pH levels (Supplementary Fig. S2B, Table S1G). This is indicative of early wound healing events occurring in epithelium^26,27^. Conversely, chronic injury resulted in the upregulation of fibrosis-related^28^ processes, such as cell migration and fiber organization (Fig. 2E, Supplementary Table S1H), alongside the downregulation of ciliogenesis (Supplementary Fig. S2C, Table S1I).

Next, a direct comparison of chronic and acute injury further unveiled transcriptional changes among the conditions (Fig. 2F, Supplementary Table S1J). In nSAECs, we noted an elevated expression of genes known to be upregulated in lung injuries, including the infection marker myoglobin (MB), which has been identified as a prognostic marker for COVID19 outcome^29,30^. Furthermore, genes associated with early infections and inflammations comprising deleted in malignant brain tumors 1 (DMBT1) and secretory genes surfactant protein A (SFTPA), secretoglobin family 3A member 1 (SCGB3A1), and SCGB2A1 were augmented in the acute compared to the chronic condition^31^. Intriguingly, these upregulated genes were implicated in pathways related to aerobic metabolism, particularly mitochondrial activity and oxidative phosphorylation in nSAECs relative to tSAECs (Fig. 2G, Supplementary Table S1K). This suggests a high cellular energy demand and aerobic turnover of lung epithelial cells to trigger physiological wound healing processes upon acute injury^32^. In contrast, a shift towards increased anaerobic, glycolytic metabolic activity has been reported in persistent, chronic injuries in lung fibrosis^33^ and was detected in the chronic injury model (Supplementary Figure S2D).

Upon chronic injury, we noticed an upregulation in the gene expression of processes involved in regulating the transcriptional machinery (Fig. 2H, Supplementary Table S1L). This suggests heightened transcriptional activity as an evolved response in tSAECs compared to nSAECs. Similar findings were reported in aberrant basal cells, a specific cell population in IPF and disease driver, which exhibit increased transcriptional activity in IPF^34^.

In summary, our findings demonstrate that these models effectively replicate the characteristic expression changes associated with early epithelial injuries as seen in mild or moderate respiratory diseases in nSAECs model, as well as advanced pro-fibrotic gene expression within tSAECs model. Notably, the initial responses in nSAECs were predominantly linked to biophysical stress responses and events related to aerobic metabolism, while these changes are further progressed into maladaptations of the transcriptional machinery in tSAECs, potentially manifesting the pro-fibrotic response.

### Fibroblasts drive the fibrotic response in chronic epithelial injury

Next, we assessed the impact of fibroblasts (LFs) on nSAECs and tSAECs. RNA-seq analysis revealed that both nSAECs and tSAECs changed their transcriptomic profiles (DEGs, p-value < 0.05) when cocultured with fibroblast (Fig. 3A, B, Supplementary Table S2A, B).

**Figure 3:**
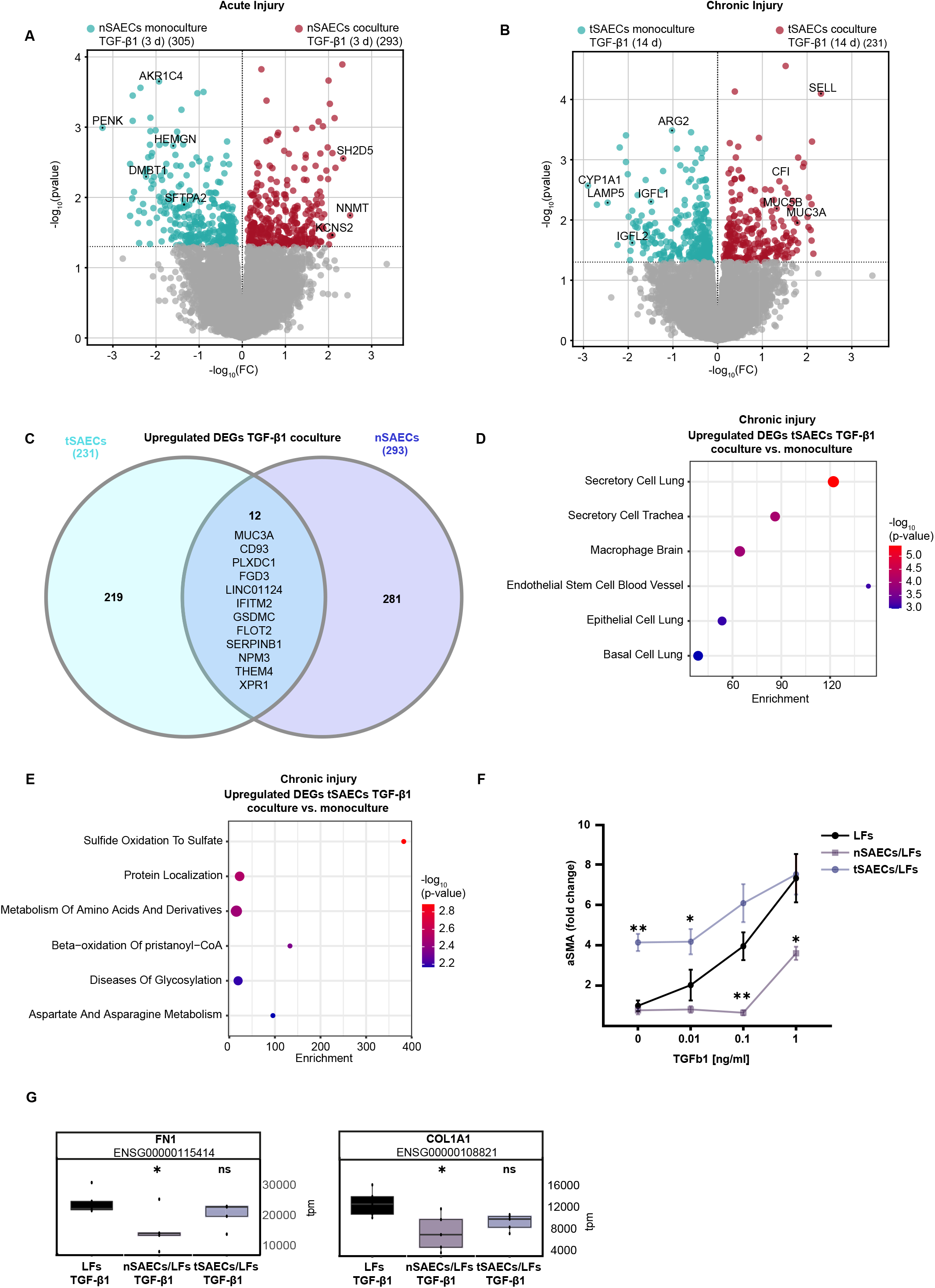
Intercellular communication of epithelial cells and fibroblasts in acute injury provokes a beneficial effect **A,** Volcano plot of all DEGs of nSAECs treated with TGF-β1 in coculture and monoculture (p-value < 0.05) shows transcriptional differences. **B**, Volcano plot of all DEGs of tSAECs treated with TGF-β1 in coculture and monoculture (p-value < 0.05) shows differences in the expression of genes. **C**, Venn diagram representing the overlap of upregulated DEGs (p < 0.05, log2FC > 0) in tSAECs and nSAECs monocultures reveals distinct responses of the cells to coculture with fibroblasts. **D**, Cell marker enrichment analysis of upregulated genes in tSAECs cocultures compared to monocultures treated with TGF-β1 shows enrichment for a secretory cell phenotype (p < 0.05, log2FC > 0). **E**, Reactome enrichment analysis of upregulated genes in tSAECs cocultures compared to monocultures treated with TGF-β1 revealed enrichment for metabolic processes involved in ECM accumulation and protein localization (p < 0.05, log2FC > 0). **F**, Alpha smooth muscle (α-SMA) expression of fibroblasts in mono- and cocultures with nSAECs and tSAECs shows attenuated effect in nSAECs. **G**, Expression levels of fibroblast-to-myofibroblast (FMT) markers COL1A1 and FN1 reveals attenuated fibroblast activation upon coculture with nSAECs.

Intriguingly, a comparison of significantly upregulated genes in nSAECs and tSAECs in cocultures revealed only a small proportion (12 genes) of shared upregulated genes (Fig. 3C), emphasizing a distinct cellular response to fibroblast crosstalk depending on the injury state. tSAEC cocultures showed an upregulation of a secretory epithelial cell phenotype compared to monocultures (Fig. 3D, Supplementary Table S2C), suggesting a functional shift in response to the chronic crosstalk. This aligns with findings in IPF, which show an increase in secreting cell phenotypes with overall mucous accumulation^20,35^. In contrast, heightened transcriptional activity was observed in monocultures, reflecting the intrinsic cellular response compared to the more communicative phenotype observed in cocultures (Supplementary Fig. S3A, Table S2D). Notably, tSAEC exhibited upregulation of genes associated with ECM-related metabolism and protein localization in coculture with LFs (Fig. 3E, Supplementary Table S2E), highlighting a change in protein function potentially linked to endoplasmatic reticulum (ER) stress and aberrant remodeling, as reported in IPF^36^. This indicates that while tSAECs alter their transcriptional program in response to intercellular communication with fibroblasts, these changes are not associated with any ameliorating responses on the chronically injured, pro-fibrotic epithelial state.

To address how acute and chronic epithelial injury alters the fibroblast phenotype in cocultures, we used increasing concentrations of TGF-β1 in tSAECs and nSAECs cocultures and monocultures of fibroblasts to investigate the fibroblast-to-myofibroblast (FMT) transition, marked by alpha smooth muscle actin (α-SMA). Chronically injured epithelial cells were able to induce FMT process in LFs even without exposure to TGF-β1, suggesting that pre-injured epithelial cells alone can activate fibroblasts (Fig. 3F). This is in line with our previous findings and further strengthens the general assumption that epithelial injury is the key initiator in the fibrotic process in IPF^7^ . Intriguingly, we observed a significantly attenuated effect of the acute treatment in coculture on the FMT process. Similar observations were noted while checking for additional FMT markers (Fig. 3G).

Taken together, our findings revealed distinct responses of acutely and chronically injured epithelial cells to LFs. While chronically injured epithelial cells activate fibroblasts and immediately induce a vulnerable, pro-fibrotic milieu, epithelial-fibroblast crosstalk in acutely injured cells promote a reparative effect.

### Intercellular communication of epithelial cells and fibroblast has a beneficial impact on acutely injured cells by inducing a metabolic turnover towards fatty acid metabolism

To further investigate the effects of intercellular communication in acute injuries, we performed proteome analysis on nSAEC mono- and cocultures (differentially expressed proteins (DEPs), p-value < 0.05, Fig. 4A). Acutely injured epithelial monocultures displayed distinct characteristics compared to other groups upon stimulation with TGF-β1, whereas the cocultures clustered together with the untreated groups (Fig. 4A, Supplementary Fig. S4A). This suggests that epithelial cells benefit from a supportive crosstalk with fibroblasts, which ameliorates the acute injury response compared to monocultures.

**Figure 4:**
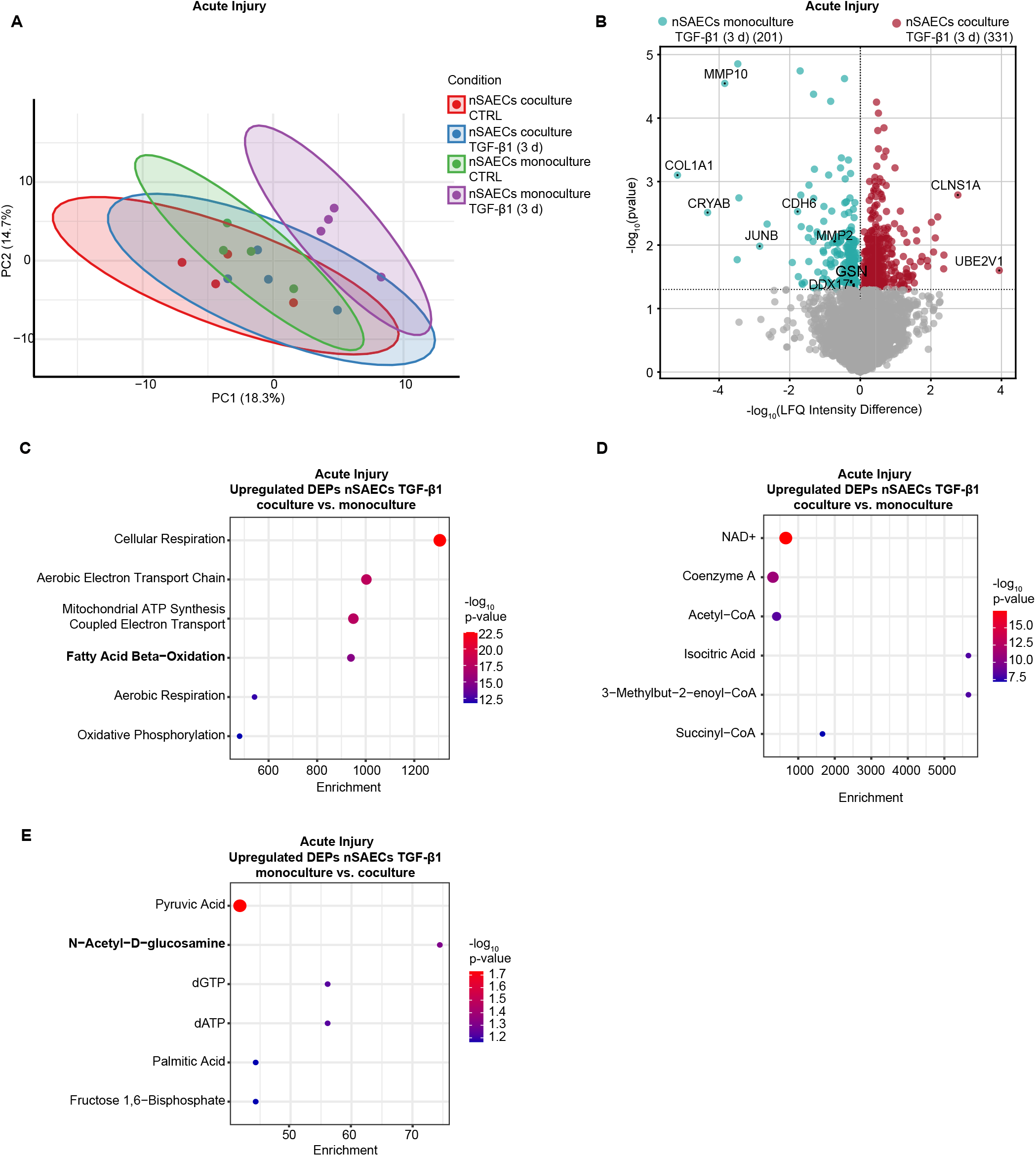
nSAECs-fibroblast coculture shows a metabolic shift. **A,** Principal component analysis (PCA) of proteomics data with top 100 variable proteins shows distinct clustering of nSAECs in mono- and cocultures and upon stimulation with TGF-β1. **B**, Volcano plot of all DEPs of nSAECs treated with TGF-β1 in coculture and monoculture (p-value < 0.05) shows differences in the proteome. **C**, GO biological process enrichment analysis of upregulated genes in nSAECs cocultures compared to monocultures treated with TGF-β1 revealed enrichment for metabolic processes especially fatty acid beta-oxidation (p < 0.05, Intensity > 0). **D**, Metabolomics Workbench Metabolite analysis shows enrichment for NAD+ in nSAECs cocultures compared to monocultures (p < 0.05, Intensity > 0). **E**, Metabolomics Workbench Metabolite analysis shows enrichment of N-Acetyl-D- glucosamine in nSAECs monocultures compared to cocultures (p < 0.05, Intensity > 0).

Next, we examined the DEPs from nSAEC cocultures and monocultures, and identified the upregulated expression of various pro-fibrotic^23^ and injury-inducing factors in monocultures (Fig. 4B, Supplementary Fig. S4B, Supplementary Table S3A), including JUNB, which has been recently reported to play an aberrant role in the development of IPF^34^. In line with this, GO term analysis of acute TGF-β1-treated nSAEC monocultures showed increased expression of proteins involved in IPF-associated tissue remodeling (Supplementary Fig. S4C, Table S3C). In contrast, reactome analysis on RNA level identified an upregulation of genes involved in gap junction assembly in TGF-β1-treated nSAEC cocultures compared to monocultures (Supplementary Fig. S4D, Supplementary Table S3D), suggesting a positive effect of fibroblasts on epithelial barrier properties within acute injuries. Intriguingly, we identified that TGF-β1-treated nSAEC cocultures exhibited high protein levels of aerobic metabolic pathways and fatty acid metabolism (Fig. 4C, Supplementary Table S3E), similar to our previous comparisons between chronic and acute injuries at the RNA level (Fig. 2D). The regenerative effects of a switch from glycolytic pathways towards fatty acid utilization have been previously reported in airway epithelial cells^37^. This is similar to findings reporting the beneficial effects of this process in wound healing^38^.

To further delineate this effect on specific metabolites, we performed an analysis on the metabolic regulation of upregulated proteins in nSAEC cocultures. Notably, we detected increased enrichment for NAD+ as a primary metabolite generated from these metabolic processes (Fig. 4D, Supplementary Table S3F), which has been frequently described as an anti-inflammatory, rejuvenating metabolite^39,40^. In contrast, monocultures favored the production of glycolytic metabolites, including N-Acetyl-D-glucosamine (GlcNAc), compared to cocultures (Fig. 4E, Supplementary Table S3G), which has recently been identified as a driver of fibrotic cell fate decision^34^.

In conclusion, we detected the beneficial effects of fibroblasts on acute injuries in epithelial cells by promoting a metabolic shift from glycolytic pathways towards unsaturated fatty acid metabolism. This shift leads to an increase in NAD+ levels and supports physiological wound healing processes. Importantly, in the acute injury model, both epithelial cells and fibroblasts collaborate to support each other and mitigate the injury-related response.

### PTX3 is an anti-fibrotic regulator and prevents pro-fibrotic response upon chronic injury

To identify key secreted proteins involved in the protective intercellular communication upon acute injury, we performed secretome analysis. Similar to the observations at the RNA and protein levels, we found that proteins, involved in aberrant tissue repair in fibrosis and lung injuries^22,41–43^, were upregulated in monocultures (differentially secreted proteins (DSP), p-value < 0.05, Fig. 5A, Supplementary Table S4A). However, certain secreted proteins which have been previously described to exert an antifibrotic role, such as CTSK, EMILIN1, and THY1, are highly secreted in cocultures^44–47^. Consistent with the RNA and protein levels, we detected an enrichment of secreted proteins associated with increased migration and injury in nSAEC monocultures compared to cocultures (Fig. 5B, Supplementary Table S4B). In contrast, the coculture secretome induced inhibition of MMPs (Fig. 5C, Supplementary Table S4C) and supported increased levels of lipid-related pathways, consistent with our previous findings (Fig. 5D, Supplementary Table S4D).

**Figure 5:**
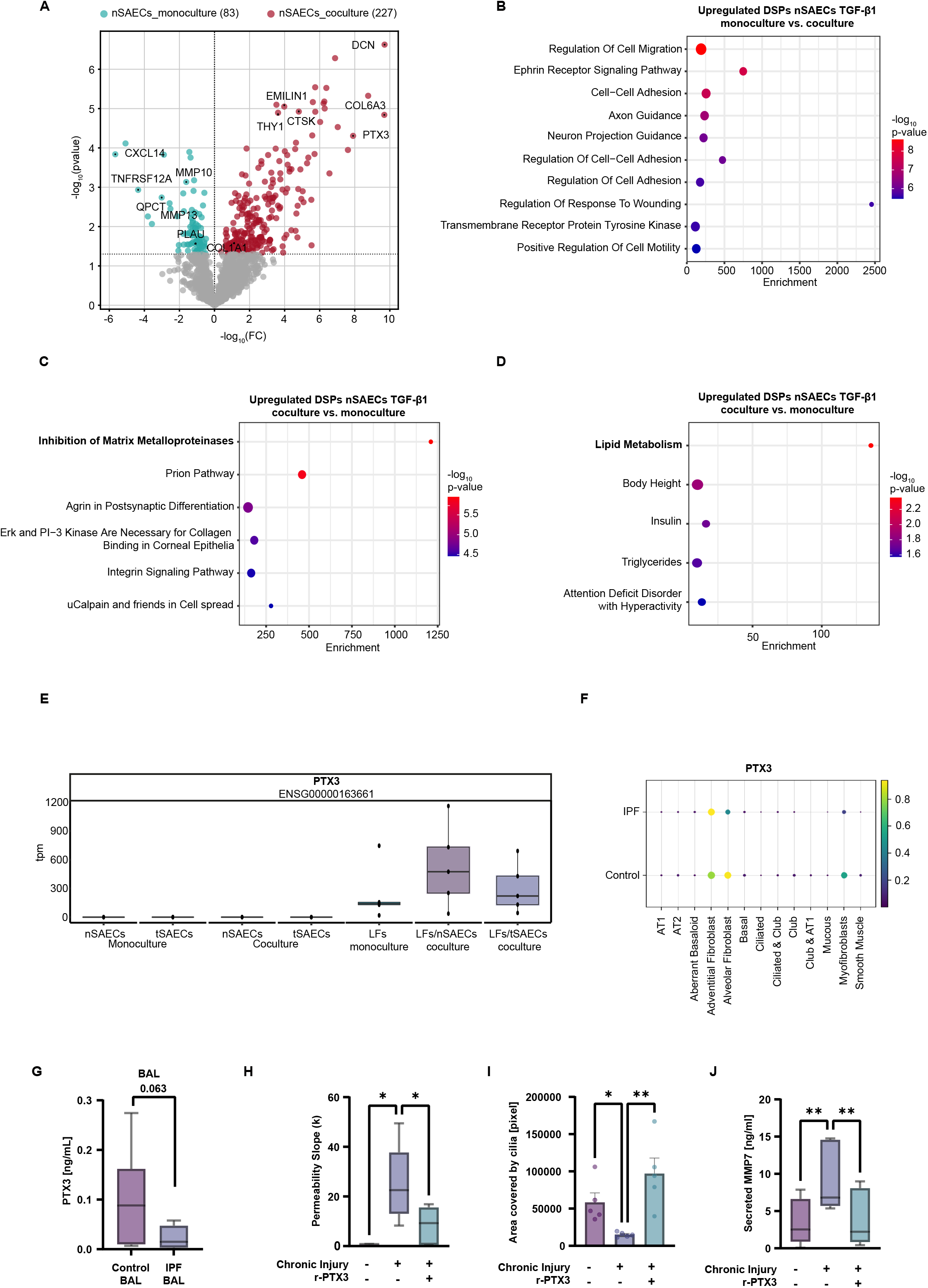
Secretome analysis revealed antifibrotic communication in nSAECs fibroblast cocultures. **A,** Volcano plot of all DSPs of nSAECs treated with TGF-β1 in coculture and monoculture (p-value < 0.05) shows differences in the secretome. **B**, GO biological process analysis shows enrichment of processes involved in tissue remodeling in nSAECs monocultures compared to cocultures (p < 0.05, Intensity > 0). **C**, KEGG analysis reveals enrichment for inhibition of MMPs in nSAEC cocultures compared to monocultures (p < 0.05, Intensity > 0). **D**, Human Database analysis shows enrichment of for lipid metabolism in nSAECs cocultures compared to monocultures (p < 0.05, Intensity > 0). **E**, TPM values of RNA-seq data shows increased PTX3 expression in acute compared to chronic injury cocultures in fibroblasts. **F**, Gene expression analysis in IPF patient scRNA-seq data showed decreased PTX3 expression in IPF fibroblasts. **G**, Human bronchoalveolar lavage (BAL) of IPF patients and control shows decrease of PTX3 in BAL samples of IPF patients (boxplot, one-sided t-test). **H**, FITC dextran permeability measurement (slope) showing a mitigation of the decreased permeability upon stimulation with PTX3 (50 ng/ml) in chronic injury (tSAEC monoculture) (boxplot, *p < 0.05, ANOVA/ Šídák’s). **I**, Area covered by ciliated cells shows increase in cilia upon stimulation with PTX3 (50 ng/ml) in tSAEC monocultures (mean +/- s.e.m., p* < 0.05, p** < 0.01, ANOVA/Holm- Šídák’s). **J**, ELISA measurement shows decrease of MMP7 secretion upon costimulation with PTX3 (50 ng/ml) (boxplot, p** < 0.01, ANOVA/Holm-Šídák’s).

**Figure 6:**
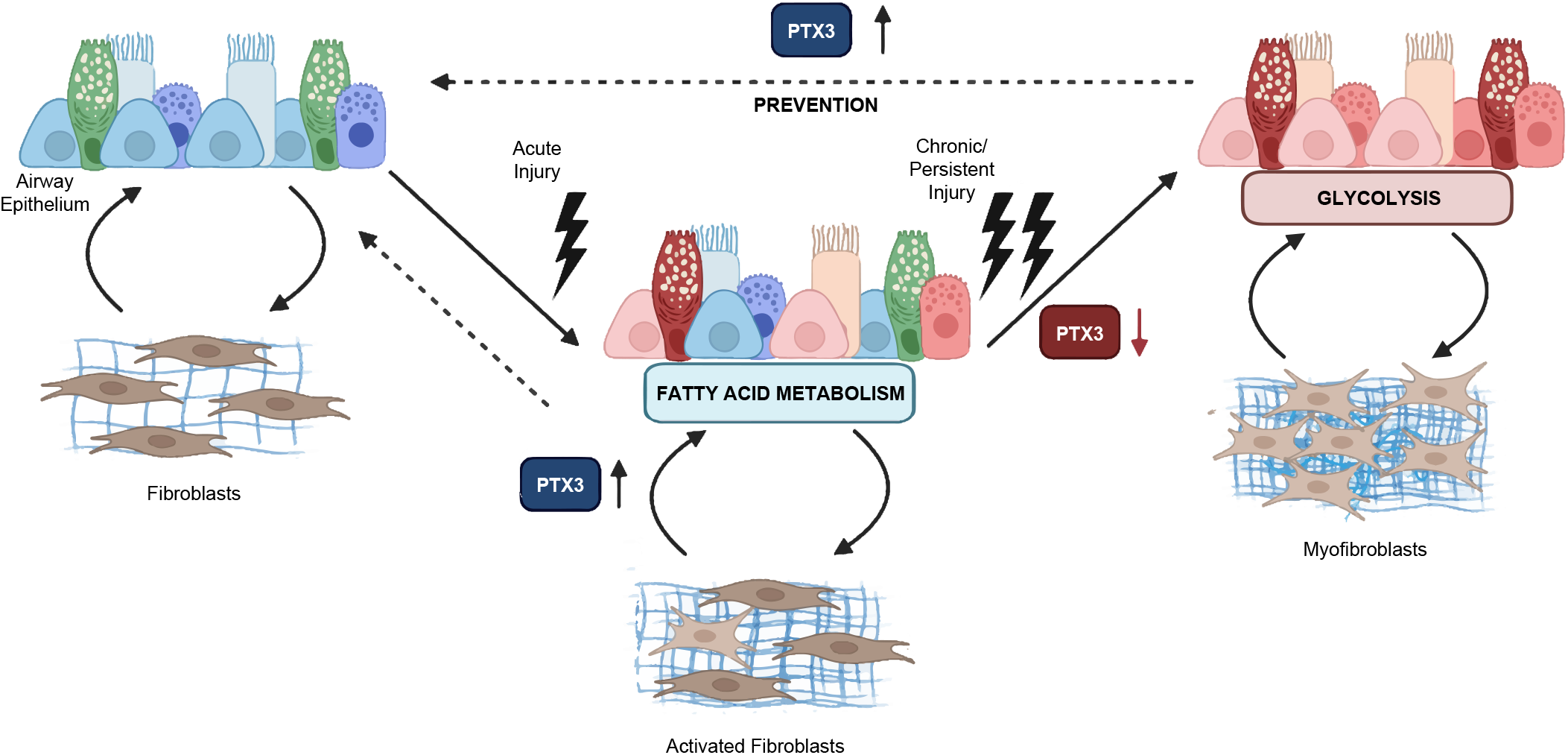
PTX3 exerts a positive role in protecting airway epithelial cells of chronic injury. **B**, Human bronchoalveolar lavage (BAL) of IPF patients and control shows decrease of PTX3 in BAL samples of IPF patients (boxplot, one-sided t-test). **C**, FITC dextran permeability measurement (slope) showing a mitigation of the decreased permeability upon stimulation with PTX3 (50 ng/ml) in chronic injury (tSAEC monoculture) (boxplot, *p < 0.05, ANOVA/ Šídák’s). **D**, Area covered by ciliated cells shows increase in cilia upon stimulation with PTX3 (50 ng/ml) in tSAEC monocultures (mean +/- s.e.m., p* < 0.05, p** < 0.01, ANOVA/Holm- Šídák’s). **E**, ELISA measurement shows decrease of MMP7 secretion upon costimulation with PTX3 (50 ng/ml) (boxplot, p** < 0.01, ANOVA/Holm-Šídák’s).

To identify specific secreted factors that are responsible for the positive effects, we compared the top ten secreted proteins in nSAEC cocultures to those in nSAEC and fibroblast monocultures. Among these proteins, only one top-secreted protein – pentraxin 3 (PTX3) - was detected (Supplementary Fig. S5A, Supplementary Table S4E). By approaching RNA data and comparing the expression levels among SAECs and fibroblast in mono-and cocultures, we observed increased expression of PTX3 in fibroblasts in acutely injured cocultures compared to chronic injury and monocultures (Fig. 5E). Moreover, expression of PTX3 was only detectable in fibroblasts, further suggesting that this protein is solely expressed and secreted by fibroblasts in this context. Next, we looked at scRNA-seq data^48^ of IPF patients and found a decreased expression of PTX3 in IPF fibroblasts compared to controls (Fig. 5F). This finding was further strengthened by analysis of human bronchoalveolar lavage (BAL) samples from IPF and control patients to assess differences in PTX3 levels. We observed a decrease in PTX3 levels within human IPF patient BAL samples (Fig. 5G), suggesting that in chronic injuries as those seen in IPF fibroblasts decrease the expression of PTX3 in the intercellular communication of epithelial cells compared to acute injuries, making them more vulnerable to the injury.

To test this hypothesis, we exposed SAECs to chronic injury in the presence of human recombinant PTX3. Intriguingly, the addition of PTX3 ameliorated the loss of epithelial barrier integrity (Fig. 5H, Supplementary Fig. S5B) and prevented the loss of ciliated cells (Fig. 5I, Supplementary Fig. S5C) upon chronic exposure to TGF-β1. Additionally, stimulation with PTX3 led to reduced levels of biomarker MMP7 (Fig. 5J, Supplementary Fig. S5D).

These findings provide novel insights into the role of PTX3 as an antifibrotic sensor on injured epithelial cells, preventing persistent injury by modifying the epithelial-fibroblast niche communications for proper repair.

## Discussion

Physiological wound healing is vital for restoring tissue homeostasis after an injury^49,50^. However, during persistent fibrosis, this tightly regulated process becomes dysregulated, leading to aberrant tissue repair^51^. In this study, we delineate the fundamental differences in acute and chronic injury concerning intercellular communication and the effect of fibroblasts on epithelial cells. Our finding reveal that fibroblast-epithelial interaction provides a protective effect in acute injuries, characterized by PTX3 secretion and a beneficial metabolic shift towards fatty acid metabolism. This ultimately facilitates the proper wound healing processes, as depicted in figure 7.

**Figure 7:** **Role of PTX3 and metabolic state in the intercellular communication of epithelial cells and fibroblasts in chronic and acute injuries.** Within acute injuries PTX3 is upregulated and secreted by fibroblasts inducing a metabolic shift of epithelial cells towards fatty acid metabolism. This mechanism is lost within the chronic injury situation. Due to the addition of PTX3 this can be restored.

While 2D air-liquid interface models of airway epithelial cells are a valuable tool for studying disease mechanisms and pharmacological testing^52^, we have established a model which not only facilitates investigations of fibroblast crosstalk, but also allows for comparisons of intercellular communication in acute and chronic injury, mimicking distinct pathological mechanisms of lung injury. Emerging evidence suggests that severe damage caused by COVID-19 may precipitate the later onset of pulmonary fibrosis^53^. Therefore, our model is relevant for studying the progression from minor lung diseases to severe outcomes.

Cells need to fine-tune certain cellular programs to adapt to alterations in their niche^1,54^. However, during chronic injuries, these programs become disrupted, leading to a loss of cellular ability to disengage and deactivate these programs. Chronic injury to epithelial cells triggers a destructive chain of events, altering niche signaling and activating fibroblasts that sustain injury programs^7,16^. This creates a self-perpetuating cycle that the cells find difficult to escape. Our findings indicate that these mechanisms are characterized by modifications in transcriptional programs within the chronic injury, rendering the cells incapable of escaping these pro-fibrotic programs. Conversely, during acute epithelial injuries, protective cellular programs are activated in response to niche signaling, with fibroblasts counteracting the pro-fibrotic response.

Cellular metabolic adaptations are a critical response to initial injury, often characterized by a shift towards anaerobic glycolytic programs^55,56^. This shift, potentially due to the rapid energy provision glycolysis offers, occurs early on, and reflects the high energy demands required for injury resolution. However, the persistence of an anaerobic metabolic state, despite its immediate benefits, is not conducive to optimal cellular function and is often indicative of a pathological condition^33,57,58^. In our acute injury model, we observed a protective effect of fibroblasts on epithelial cells, mediated through a metabolic shift towards fatty acid metabolism. This shift, which induces NAD+ may represent a key mechanism by which cells escape the detrimental effects of injury. In contrast, in chronic conditions such as IPF, glycolytic activity remains prevalent^33,59^. We suggest that this persistent glycolytic activity could be a contributing factor to the pathogenesis of IPF. However, further investigations are needed to confirm this hypothesis and to elucidate the potential therapeutic implications of metabolic shifts in lung injury and repair.

Restoring physiological conditions through niche modifications could initiate proper wound healing in chronic injury. Secretome analysis identified PTX3, a protein secreted by fibroblasts, as a key player in protective injury response and repair. We found that addition of recombinant PTX3 can protect chronically injured epithelial cells from a fibrotic fate. We suggest that this effect is caused by either binding of PTX3 to epithelial cell receptors and initiation of intercellular, injury-resolving programs, or the neutralization of pro-fibrotic factors to ultimately escape the perpetuating fibrotic cell fate as described in other findings^60,61^ . This effect potentially activates a shift in the metabolic state of epithelial cells^62,63^ ; however, the role of PTX3 in metabolic regulation requires further elucidation.

In summary, our study illuminates the fundamental differences in acute and chronic injury with respect to intercellular communication and highlights the protective effect of fibroblasts on epithelial cells in acute injuries. Our findings underscore the importance of understanding the interplay between cellular metabolic adaptations and injury response and suggest that a shift towards fatty acid metabolism may offer a protective mechanism against acute injury. Moreover, we propose PTX3 as novel regulator positively impacting epithelial injuries. This research provides valuable insights into the dysregulated intercellular communication in IPF and lays the groundwork for future investigations into therapeutic interventions that could leverage the protective effects of PTX3.

## MATERIALS AND METHODS

### Cultivation of primary cells

Primary healthy human lung airway epithelial cells (SAECs, Lonza, #CC-2547) were used in passages 2-3. They were expanded in flasks on type I collagen (Corning, #354236) pre-coated cell culture flasks in PneumaCult™-Ex Plus Medium (STEMCELL Technologies, #05040) until confluency of around 80 %. For air-liquid-interface (ALI) cultivation and differentiation, 30 000 cells/well SAECs were seeded on collagen-coated transwells and grown until confluency in PneumaCult™-Ex Plus Medium. Airlift was conducted by changing medium to PneumaCult™-ALI Medium (STEMCELL Technologies, #05001) and exposing cells apically to air.

Healthy human lung fibroblasts (LFs) (Lonza, #CC-2512) were used in passages 4-7. They were expanded in flasks in Fibroblast Basal Medium (Lonza, #CC-3131) with FGM2- Fibroblast Growth Medium BulletKit (Lonza, CC-3132) until confluency of around 70 %.

### Epithelial-mesenchymal coculture

SAECs were differentiated in air liquid interface (ALI) on transwells for 10 days until differentiated cells were initially observed. Afterwards, ALI SAECs were treated for 14 days with 1 ng/ml TGF-β1 (Biotechne, #240-B-002) to induce chronic injury on SAECs. Additionally, 10 ng/ml or 50 ng/ml recombinant human PTX3 (Biotechne, #10292-TS-050) was used in combination with TGF-β1 and vehicle-treated cells implied as controls. For the acute injury model, ALI SAECs were cultivated for 14 days without any stimuli.

After 14 days, both, SAECs for chronic and acute injury were transferred into cocultures with human lung fibroblasts +/- 1 ng/ml TGF-β1 and +/- PTX3 in the basolateral chamber for additional 3 days. Before setting up the coculture, both, SAECs and LFs were starved for 6-8 h in normal DMEM (Gibco^TM^, # 31966021), which was then further used as coculture medium.

### Hematoxilin eosin staining

Chronically injured SAECs and respective untreated controls were seeded and cultivated as previously described and afterwards fixed in 4 % paraformaldehyde (PFA) for 20 min at room temperature (RT). Whole transwell membranes were cut out of the inserts and embedded in HistoGel^TM^ (Richard-Allan Scientific, #HG-4000-012) and transferred to a paraffin cassette followed by incubation for 30 min at 4 °C. The cassette was embedded in paraffin and 3 µm slices were cut by using microtome. For Hematoxilin-Eosin staining, the respective sections were deparaffinized and stained according to standard protocols. Images were taken using a Zeiss Axio Observer system (Zeiss, Oberkochen, Germany) with 20× objective of an Axio Imager Z1 (Zeiss, Oberkochen, Germany).

### FITC dextran permeability assay

For permeability measurement using 10 kDa fluorescein isothiocyanate (FITC) dextrane (Sigma Aldrich, #FD10S), the medium of the basolateral chamber of the respective SAECs conditions was exchanged to RPMI without phenol red (GibcoTM, #11835030). 5 mg/ml FITC dextrane in RPMI was added to the apical chamber and samples from the basolateral chamber were taken after 0 min, 30 min, 60 min, and 90 min. Relative fluorescence intensity (excitation 490 nm, emission 520 nm) was measured using the SpectraMax M5 (Molecular Devices, San Jose, CA, US). Slopes were calculated by simple linear regression.

### Cilia beating measurement

Active cilia and cilia beat frequency were assessed by image stacks of 2D+time as described in previous work^64^. Briefly, four different regions of the transwells were imaged using 32x objective of an Axiovert25 (Zeiss, Oberkochen, Germany) and an acA 1300-200 µm black and white USA-3.0 high speed camera (Basler, Ahrensburg, Germany). 100 frames per second for a total of 6 seconds were captured to record cilia movement. HALCON 13.0.2 software toolbox (MVTec Software, Munich, Germany) was used to develop applications for image capture and analysis and Analyze (AnalyzeDirect, Overland Park, KS, US) was utilized to visualize the image stack. Area covered by cilia was determined by quantifying detectably moving pixels within the chosen area. Ciliary beating frequency was calculated by the average frequency of the change in grey values.

### Nanoindentation

Stiffness measurement of SAECs on transwells was performed via interferometry-based optical fibre-top nanoindentation by Pavone (Optics11Life, Amsterdam, Netherlands). Cantilevers with stiffnesses of 0.029 N/m and 0.017 N/m were employed and an area of 32.5 µm^2^ for measurements used. Before the first measurements, a matrix scan was conducted where nanoindentation of 35 points per well was performed.

### ELISA

Supernatants were centrifuged and diluted 1:10-1:100. ELISA measurements for MMP7 (Biotechne, #DY907), MMP10 (Biotechne, #DY910), and PTX3 (Thermo Fisher, #EH386RB) were conducted according to manufacturer’s instructions. Relative fluorescence intensity (excitation 450 nm, emission 540 nm) was measured via SpectraMax M5 and SoftMax® Pro software (Molecular Devices, San Jose, CA, US). Total protein concentration was assessed by standard curve.

### RNA extraction and RNA sequencing

Chronically injured and acutely injured SAECs were cultivated as previously described and lysed in RLT plus buffer (Qiagen, #1053393). Total RNA was isolated and purified using the MagMAX^TM^ 96 Total RNA Isolation Kit (Thermo Fisher, #AM1830) following manufacturer’s instruction. Total RNA was assessed for quantity using the fluorescence-based Broad Range Quant-iT RNA Assay Kit (Thermo Fisher, # Q10213) and quality using the Standard Sensitivity RNA Analysis DNF-471 Kit (Agilent, # DNF-471-0500) on a 48-channel Fragment Analyzer (Agilent, Santa Clara, CA, US). All samples had RIN values > 9.

Samples were normalized on the MicroLab STAR automated liquid platform (Hamilton, Bonaduz, Switzerland). A total of 200 ng RNA was used for library construction using the NEBNext Ultra II Directional RNA Library Prep Kit for Illumina (New England Biolabs, #E7760), together with the NEBNext Poly(A) mRNA Magnetic Isolation Module (New England Biolabs, #E7490) and NEBNext Multiplex Oligos for Illumina (New England Biolabs, #E7600). RNA-seq libraries were processed on a Biomek i7 Hybrid instrument (Beckman Coulter, Brea, CA, US) using 12 PCR cylces following manufacturer’s instruction (except for the use of Ampure XP beads (Beckman Coulter, #A63880) instead of SRPIselect beads). Final RNA-seq libraries were then quantified by the High Sensitivity dsDNA Quanti- iT Assay Kit (Thermo Fisher, #Q33120) on a BioTek Synergy HTX (Agilent, Santa Clara, CA, US). And assessed for their size distribution and adapter dimer presence using the High Sensitivity Small Fragment DNF-477 Kit (Agilent, # DNF-477-0500) on a 48-channel Fragment Analyzer (Agilent, Santa Clara, CA, US). All sequencing libraries were then normalized on the MicroLab STAR and pooled. Additionally, PhiX Control v3 (Illumina, #FC-110-3001) was spiked in. The pooled libraries were clustered on a Paired-End Flow Cell and sequenced on a HiSeq 4000 Sequencing System (Illumina, San Diego, CA, US) with dual index, paired-end reads at 75 bp length. Read parameters: Rd1: 76 bp, Rd2: 8 bp, Rd3: 8 bp, Rd4: 76 bp with an average sequencing depth of around 30 million pass-filter reads per sample.

### RNA-seq analysis

Demultiplexing was conducted using bcl2fastq2 v2.20.0.422 from Illumina. Other pre- processing was performed as previously described^65^. Differentially expressed gene (DEG) analysis was carried out on the mapped counts derived from featureCount using the limma/voom R-package with Benjamini–Hochberg correction^66,67^. Genes with a p-value < 0.05 were considered differentially expressed. DEGs were further analysed using EnrichR^68^ and visually represented using SRplot^69^.

### Sample preparation for LC-MS/MS

Supernatant samples were buffer exchanged using 3.5-kDa Amicon filter (Millipore) to 8 M urea, 50 mM ammonium bicarbonate buffer. While in the filter, proteins were reduced with 5 mM tris(2-carboxyethyl)phosphine (TCEP) at 30°C for 60 min, and subsequently alkylated with 15 mM iodoacetamide (IAA) in the dark at room temperature for 30 min. The buffer was then exchanged again to 1 M urea, 50 mM ammonium bicarbonate, the sample was recovered from the Amicon tube into a new microfuge tube and protein concentration was determined using bicinchoninic acid (BCA) protein assay (Thermo Scientific). Proteins were subjected to overnight digestion with mass spec grade Trypsin/Lys-C mix (Promega). Following digestion, samples were acidified with formic acid (FA) and subsequently peptides were desalted using AssayMap C18 cartridges mounted on an AssayMap Bravo Platform (Agilent Technologies).

Cells were lysed in 8M urea, 50 mM ammonium bicarbonate (ABC) and Benzonase, and the lysate was centrifuged at 14,000 × g for 15 minutes to remove cellular debris. Supernatant protein concentration was determined using a bicinchoninic acid (BCA) protein assay (Thermo Scientific). Disulfide bridges were reduced with 5 mM tris(2- carboxyethyl)phosphine (TCEP) at 30°C for 60 min, and cysteines were subsequently alkylated with 15 mM iodoacetamide (IAA) in the dark at room temperature for 30 min. Urea was then diluted to 1 M urea using 50 mM ABC, and proteins were subjected to overnight digestion with mass spec grade Trypsin/Lys-C mix (Promega). Following digestion, samples were acidified with formic acid (FA) and subsequently peptides were desalted using AssayMap C18 cartridges mounted on an AssayMap Bravo Platform (Agilent Technologies).

### LC-MS/MS analysis

Prior to LC-MS/MS analysis, dried peptides were reconstituted with 2% ACN, 0.1% FA and concentration was determined using a NanoDrop^TM^ spectrophometer (ThermoFisher). Samples were then analyzed by LC-MS/MS using a Proxeon EASY-nanoLC system (ThermoFisher) coupled to an Orbitrap Fusion Lumos mass spectrometer (Thermo Fisher Scientific). Peptides were separated using an analytical C18 Acclaim PepMap column (75µm x 500 mm, 2µm particles; Thermo Scientific) at a flow rate of 300 nL/min (60°C) using a 75-min gradient: 1% to 5% B in 1 min, 6% to 23% B in 44 min, 23% to 34% B in 28 min, and 34% to 48% B in 2 min (A= FA 0.1%; B=80% ACN: 0.1% FA). The mass spectrometer was operated in positive data-dependent acquisition mode. MS1 spectra were measured in the Orbitrap in a mass-to-charge (*m/z*) of 375 – 1500 with a resolution of 60,000 at *m/z* 200. Automatic gain control target was set to 4 x 10^5^ with a maximum injection time of 50 ms. The instrument was set to run in top speed mode with 2-second cycles for the survey and the MS/MS scans. After a survey scan, the most abundant precursors (with charge state between +2 and +7) were isolated in the quadrupole with an isolation window of 1.6 *m/z* and fragmented with HCD at 30% normalized collision energy. Fragmented precursors were detected in the ion trap as rapid scan mode with automatic gain control target set to 1 x 10^4^ and a maximum injection time set at 35 ms. The dynamic exclusion was set to 20 seconds with a 10 ppm mass tolerance around the precursor.

### Mass spectrometry and data processing

All raw files of secretomics and proteomics were analyzed using MaxQuant quantitative proteomics software (version 1.5.5.1) with the integrated Andromeda Search engine^70^ against a target-decoy version of the curated human UniProt proteome (downloaded January 2019), excluding isoforms, as well as the GPM CRAP sequence. The mass tolerance for the first search was set to 20 ppm, while for the main search, the tolerance was narrowed to 4.5 ppm. The fragment ion mass tolerance was set to 20 ppm. Trypsin was specified as digestion enzyme in specific cleavage mode, allowing up to two missed cleavages. Carbamidomethylation of cysteine was applied as fixed modification and protein N-terminal acetylation and methionine were specified as variable modifications. A 1 % target-decoy-based false discovery rate (FDR) was applied for spectrum and protein identification.

### Mass spectrometry data analysis

Quantitative analysis of the processed proteome and secretome data was conducted using Perseus (version 2.0.9.0). Initially, potential contaminants, proteins identified from the reverse database, and proteins identified only by site were excluded from the analysis. Proteins were further filtered to retain those with valid intensity values in at least 70 % of samples in at least one group and with intensities greater than 0. Intensity values were log2-transformed and missing values were replaced from normal distribution. Welch’s t-test was applied to identify differentially expressed proteins between experimental groups. Differentially expressed proteins with p-values < 0.05 were considered significant. SR plot^69^ and R (version 4.4.2, R studio version 2024.09.0+357) were used for data visualization and correlation calculations using pearson coefficient.

### Statistical analysis and reproducibility

GraphPad prism (version 10.1.2) was utilized to perform statistical analysis. To determine significance the following test were applied: t-test, nonparametric t-test with Mann Whitney test, one-way ANOVA. Gaussian distribution was assessed for all tests requiring normal distribution with the Kolmogorov-Smirnov test and Shapiro-Wilk test (α = 0.05). For all experiments at least three biological replicates were implied.

## Acknowledgements

We thank J. Garnett for initiating the first experiments, F. Ramirez for assistance and support with RNA-seq, F. Herrmann for provision of primary fibroblasts, the Histology Lab of the Drug Discovery Science Team and V. Schröder for technical assistance and support. This work was funded by Boehringer Ingelheim Pharma GmbH und Co. KG. All drawings were created using Biorender.com.

## Author contributions

MB conceptualized, designed, performed, and supervised most of the experiments and analyzed data. HQL, CV and IK performed sequencing experiments. MB and FR analyzed sequencing data. MB analyzed proteomics and secretomics data. IK, JH, EP performed experiments. MJT, ML, HS, and FG provided conceptual advice. HQL conceived, designed experiments, and supervised the study. MB and HQL wrote the paper. All authors commented on and edited the manuscript.

## Data availability

RNA-seq data or proteomics/secretomics generated within this study will be deposited in the Gene Expression Omnibus (GEO) or MASSIVE and will be made accessible upon acceptance of the manuscript. All other data is available on request.

---

**Supplementary Figure S1: Chronic injury induces pro-fibrotic response in epithelial cells.**

**A,** Cilia beating frequency measurement shows no change in frequency in control and chronic injured epithelial cells.

**B**, ELISA measurement shows significant increase of MMP10 in tSAECs (boxplot, p* < 0.05, t-test).

**Supplementary Figure S2: Acute and chronic injury differ in their transcpritional response.**

**A,** GWAS analysis reveals enrichment for COVID-19 in nSAECs monocultures (p < 0.05, log2FC > 0).

**B**, GO biological process analysis reveals upregulation of cellular stress responses in nSAEC monocultures (p < 0.05, log2FC > 0).

**C**, GO biological process analysis shows enrichment for cilium organization in tSAEC monocultures (p < 0.05, log2FC > 0).

**D**, KEGG analysis reveals enrichment for glycolysis in tSAECs monocultures treated with TGF-β1 (p < 0.05, log2FC > 0).

**Supplementary Figure S3: Epithelial-fibroblast coculture changes the transcriptome of nSAECs and tSAECs.**

**A,** GO biological process analysis shows enrichment for increased transcriptional activity (p < 0.05, log2FC > 0).

**Supplementary Figure S4: Epithelial-fibroblast crosstalk is protective.**

**A**, Hierarchical clustering of the top 50 differentially expressed proteins (DEPs) from proteomics analysis in nSAECs monocultures treated with TGF-β1 compared to control and coculture conditions shows distinct clustering (p < 0.05).

**B**, GSEA reflects weak enrichment of an IPF lung gene signature in nSAEC monocultures compared to nSAEC cocultures (p < 0.05).

**C,** Reactome analysis shows increase in tissue remodeling processes in nSAECs monocultures vs. cocultures (p < 0.05, Intensity > 0).

**D**, Reactome analysis of upregulated genes in nSAECs cocultures treated with TGF-β1 shows enrichment for gap junction assembly (p < 0.05, log2FC > 0).

**Supplementary Figure S5: PTX3 ameliorates the pro-fibrotic response in chronically injured epithelial cells.**

**A,** Venn diagram representing the overlap of top ten upregulated DSPs in nSAEC cocultures compared to monocultures and LF monocultures compared to nSAEC cocultures (p < 0.05, Intensity > 0) reveals PTX3 as top hit.

**B**, FITC dextran permeability measurement (slope) showing a mitigation of the decreased permeability upon stimulation with PTX3 (10 ng/ml) in chronic injury (tSAEC monoculture) (boxplot, *p < 0.05, ANOVA/ Šídák’s).

**C**, Area covered by ciliated cells shows increase in cilia upon stimulation with PTX3 (10 ng/ml) in tSAEC monocultures (mean +/- s.e.m., p* < 0.05, p** < 0.01, ANOVA/Holm-Šídák’s).

**D**, ELISA measurement shows decrease of MMP7 secretion upon co-stimulation with PTX3 (10 ng/ml) (boxplot, p** < 0.01, ANOVA/Holm-Šídák’s).

