## Supplementary figures and images for "PTX3 Governs Fibroblast-Epithelial Dynamics in Lung Injury and Repair"

### Supplementary Figure 1

**Bammert et al. 2024 - Supplementary Figure 1**

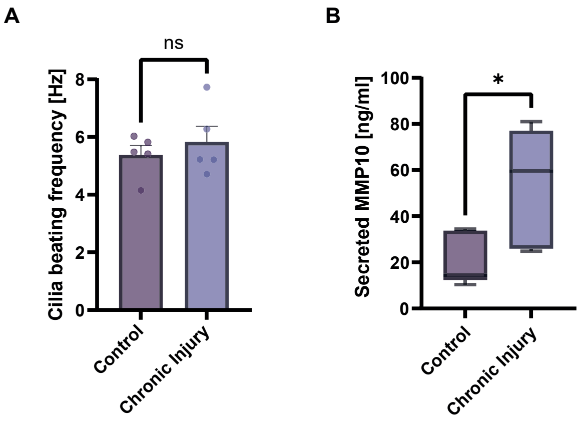

### Supplementary Figure 2

#### Bammert et al. 2024 - Supplementary Figure 2

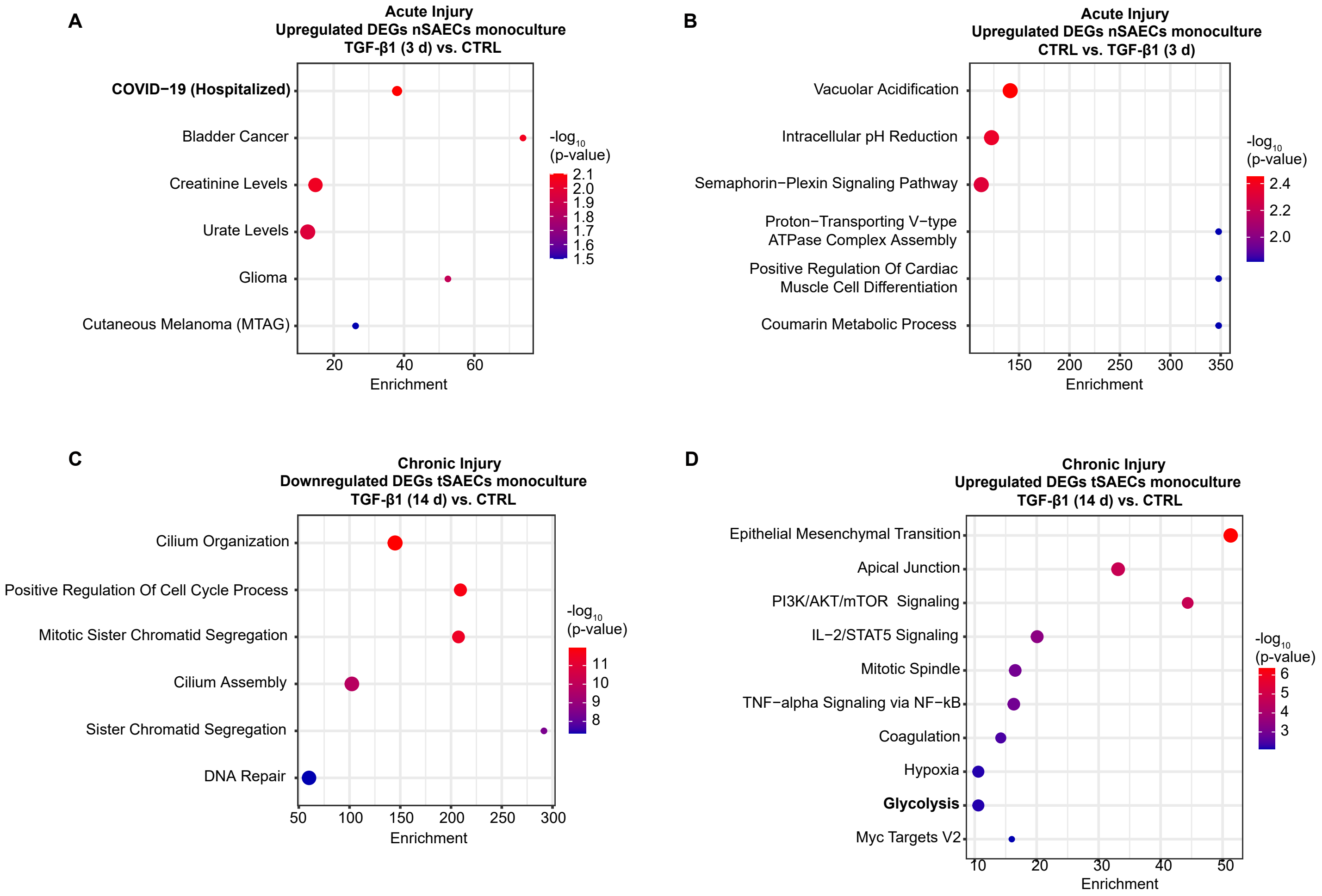

### Supplementary Figure 3

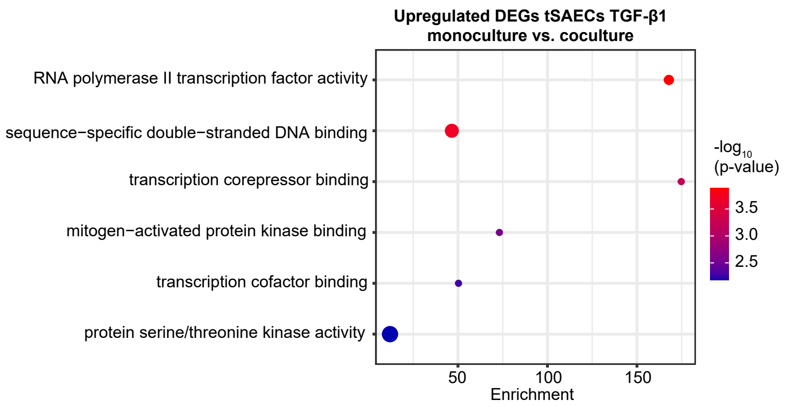

### Supplementary Figure 4

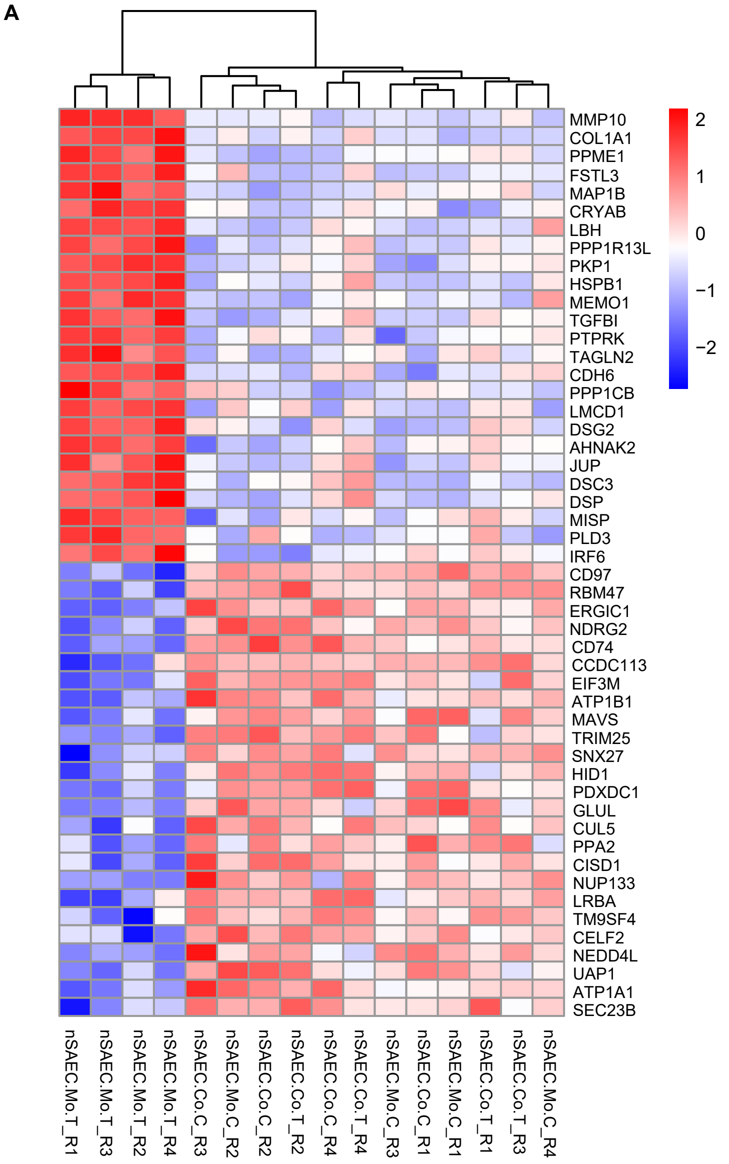

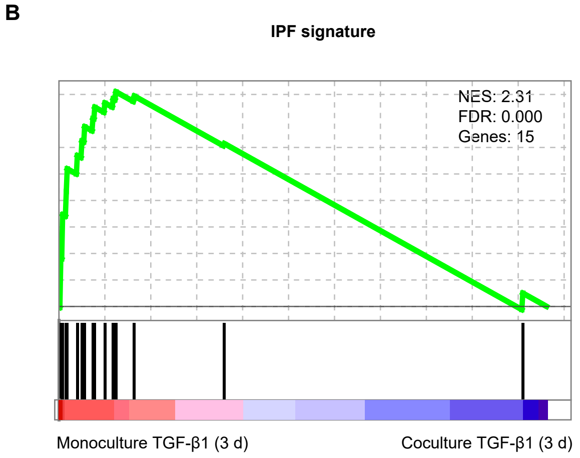

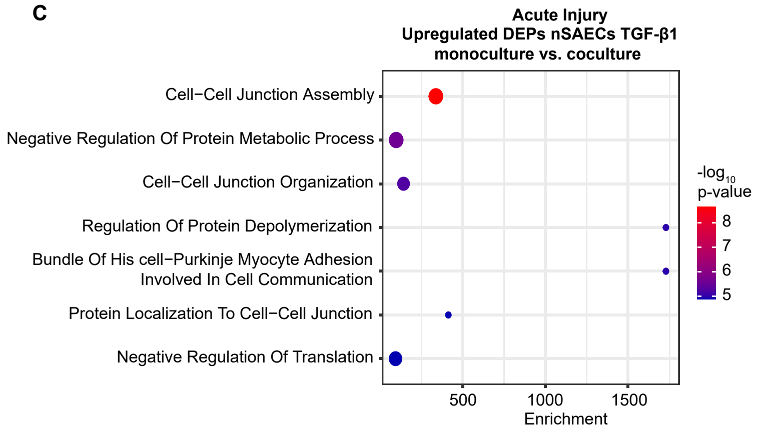

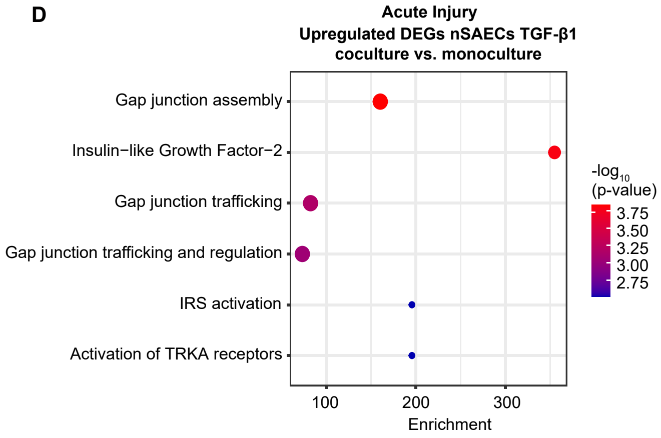

### Supplementary Figure 5

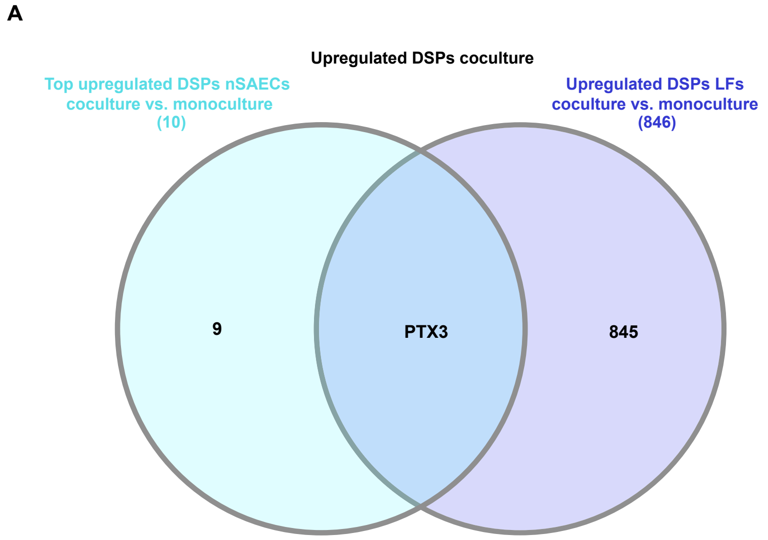

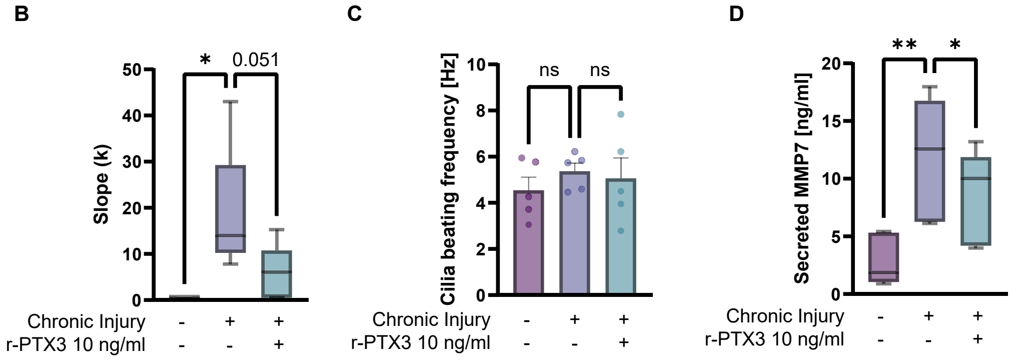
